# Analysis of polyclonal vector integration sites using Nanopore sequencing as a scalable, cost-effective platform

**DOI:** 10.1101/833897

**Authors:** Ping Zhang, Devika Ganesamoorthy, Son Hoang Nguyen, Raymond Au, Lachlan J. Coin, Siok-Keen Tey

## Abstract

Vector integration site analysis can be important in the follow-up of patients who received gene-modified cells, but current platforms based on next-generation sequencing are expensive and relatively inaccessible. We analyzed polyclonal T cells transduced by a gammaretroviral vector, SFG.iCasp9.2A.ΔCD19, from a clinical trial. Following restriction enzyme digestion, the unknown flanking genomic sequences were amplified by inverse polymerase chain reaction (PCR) or cassette ligation PCR. Nanopore sequencing could identify thousands of unique integration sites within polyclonal samples, with cassette ligation PCR showing less bias. The assay is scalable and requires minimum capital, which together enable cost-effective and timely analysis.

## Introduction

Gene modification can be highly efficient and effective in conferring specific biological traits to a cellular therapeutic. In a majority of cases, gene modification involves the integration of one or more copies of a transgene into the host cell genome, which is passed down to all its progenies. Although targeted transgene integration using CRISPR/cas9 and other genome editing techniques hold great promise and may well be the path of the future [1, 2], the vast majority of current gene-modified cellular therapeutics use gammaretroviral, lentiviral or non-viral vectors that are non-targeted and can integrate at multiple sites, with some predilection for open chromatin and transcriptionally active regions [3–6]. Analysis of vector integration sites can provide critical information on the clonality of gene-modified cells and potential biological impacts of specific transgene insertion sites, including the potential for insertional mutagenesis through the inactivation of tumor suppressor genes or activation of proto-oncogenes, such as *LMO2* [7, 8] and *EVI1* [9]; or alternatively, enhanced therapeutic efficacy, such as enhanced chimeric antigen receptor (CAR) T cell function through transgene disruption of *TET2* [10].

In general, the analysis of transgene integration sites involves polymerase chain reaction (PCR) amplification of the flanking genomic sequences, followed by sequencing of the PCR amplicons. This can be achieved by a number of methods, with one of the most commonly used being linear-amplification mediated PCR (LAM-PCR) or similar methodology followed by short-read high-throughput sequencing on Illumina or similar platforms [3, 11, 12]. These next generation sequencing platforms can generate a very large number of high-quality reads per flow cell, and thus provide economy of scale. However, the flow cells are expensive, the cost per run is high, and it is therefore necessary to pool a large number of samples to be cost efficient. In addition, short read lengths can result in inefficient genome alignment. Oxford Nanopore sequencing is a relatively new sequencing platform which directly sequences a strand of DNA as it passes through a Nanopore [13, 14], and is capable of ultra-long reads, as long as 2 megabases. Additionally, the sequencing flow cells from Oxford Nanopore Technologies are relatively inexpensive and sequencing can be performed in any laboratory without the need for dedicated sequencing equipment. Hence, it may be a cost-efficient platform for integration site analysis for smaller cell therapy centers where there are low sample volumes and potentially limited access to high-cost sequencing instruments.

We recently conducted a phase I clinical trial using T cells that were transduced with a gammaretroviral vector that carried the inducible caspase 9 *(iCasp9)* safety switch [15]. In order to understand the *in vivo* clonal dynamics of the infused T cells, we developed a method to analyze the vector integration sites in patient samples using inverse PCR followed by Nanopore sequencing [15, 16]. In this paper, we describe in detail the inverse PCR methodology and an improved methodology using cassette ligation PCR, which has less bias, both followed by Nanopore sequencing. We show that Nanopore sequencing can be a readily accessible platform for vector integration site analysis. It has some limitations in regard to read quality and single nucletoide resolution, both of which are partially offset by the longer read lengths and are likely to improve with newer generations of Nanopore sequencers and refinement in methodology.

## Materials and Methods

### Retroviral vector, patient samples and generation of transduced Jurkat cell clones

The gammaretroviral vector, SFG.iCasp9.2A.ΔCD19, has been previously described [17]. It encodes a safety switch, inducible caspase 9 *(iCasp9)*, and truncated CD19 (ΔCD19), which enables the detection of transduced cells by flow cytometry. The vector was pseudotyped with Gibbon ape leukemia virus (GALV) envelope [17]. Clinical samples were obtained from a phase I clinical trial using donor-derived *iCasp9*-transduced T cell addback as previously reported [15]. The infused cell product and peripheral blood mononuclear cells (PBMC) from Patient #1 at days 369 and 1332 after *iCasp9*-transduced T cell addback were analyzed. To generate cell lines with clonal viral integrants, Jurkat cells (2×10^5^ cells in 0.5mL) were added to 1.5mL of retroviral supernatant on retronectin-coated non-tissue culture-treated 24-well plate, centrifuged at 1000g, 32°C for 40 minutes, and incubated at 37°C, 5% CO_2_ for 24 hours. The transduced Jurkat cells were single cell cloned by limiting dilution in 96-well flat-bottom plates and clones from the lowest dilution were expanded.

### DNA extraction and restriction enzyme digestion

Genomic DNA was extracted using Purelink™ genomic DNA mini kit (Invitrogen, Waltham, MA) according to the manufacturer’s instructions. Genomic DNA was digested with two 6-cutter restriction enzymes which generate compatible cohesive ends (underlined): *NcoI*, which cuts at C/<u>CATG</u>G and *BspHI*, which cuts at T/<u>CATG</u>A (both from New England BioLabs, Ipswich, MA). *NcoI* has 3 cut sites and *BspHI* has 1 cut site within the transgene, with the most distal cut site generated by *NcoI* at 1185 base pair (bp) from the junction between the vector insert and the flanking genomic DNA (Fig.1A). Note that these restriction enzymes do not have any cut sites within the 5’ and 3’ long terminal repeats (LTR), which flank the transgene and are identical in sequence and orientation. This assay design avoids restriction enzyme digestion within the 5’LTR, which can result in the downstream amplification of the internal transgene sequence, producing non-informative reads. The resulting DNA fragments were circularized for inverse PCR or ligated to linker cassettes for downstream PCR amplification.

**Figure 1.**
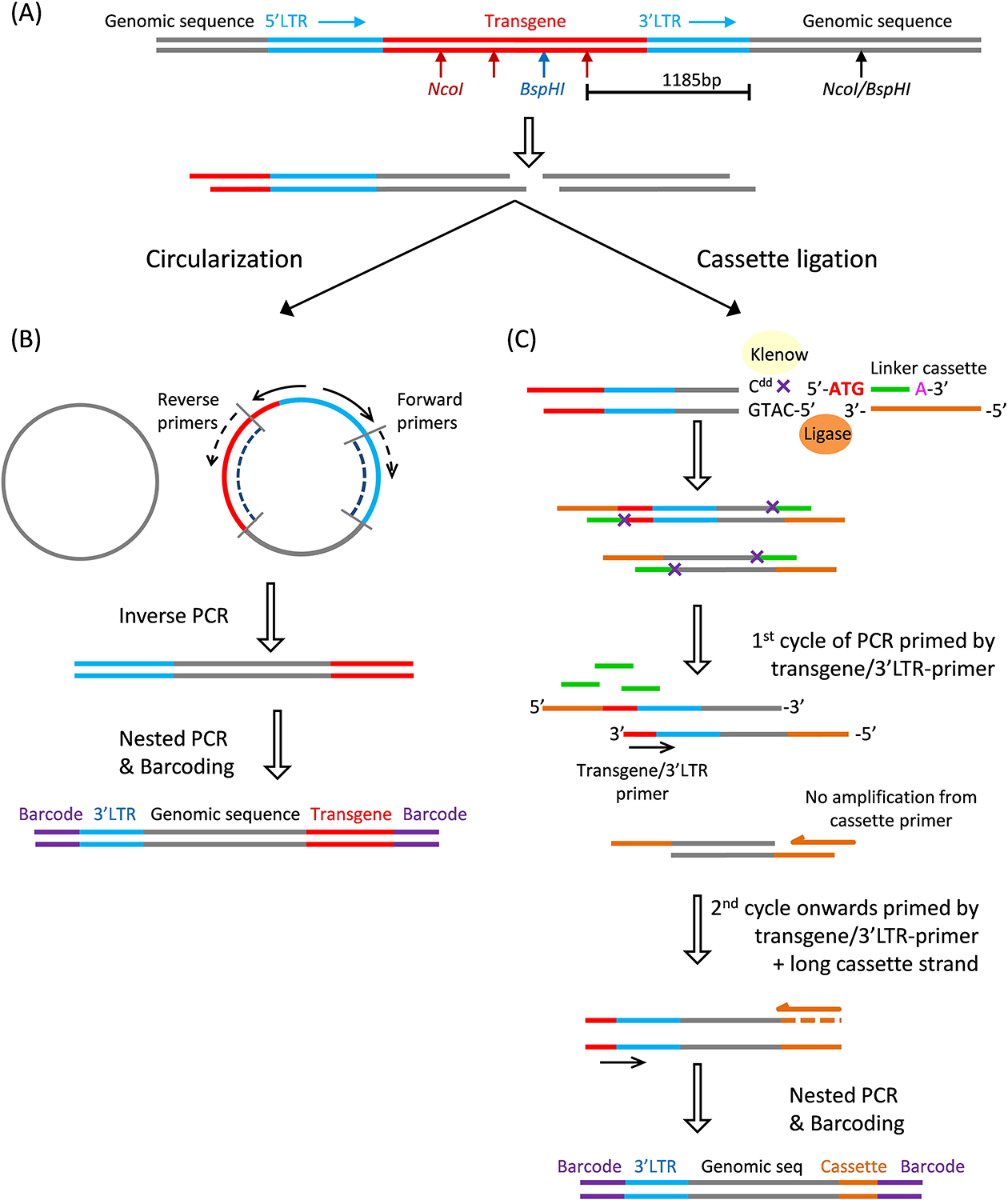
Schematic for PCR amplification of flanking genomic sequences. (A) Genomic DNA is digested with two 6-cutter restriction enzymes, *NcoI* and *BspHI*, which together are anticipated to cut at approximately 2 kb intervals. There are 4 restriction sites within the transgene sequence, the most distal of which is 1185 bp from the 3’LTR / genomic junction. *NcoI* and *BspHI* generate identical 4-nucleotide 5’ overhangs: 5’-CATG-3’, which can be circularized for inverse PCR or ligated to linker cassettes. (B) Inverse PCR begins with circularization with T4 DNA ligase, followed by PCR amplification of the unknown flanking genomic sequences using primers targeting the 3’LTR and the 3’LTR/distal transgene junction. This is followed by nested PCR, which incorporates tailing sequences for subsequent barcoding. The combined lengths of the dotted lines in the inner circle indicate the minimum theoretical length prior to the addition of tailing sequences and barcodes. (C) The ligation cassette comprises two partially complementary strands: a 27-nucleotide strand and a 14-nucleotide strand, the latter with a mismatched A at the 3’ end and a 5’overhang (5’-ATG-3’). Before cassette ligation, the genomic DNA fragments are filled with a single ddCTP to prevent elongation or ligation at the recessed 3’ end. Cassette ligation results in a nick on this strand, indicated by ‘X’. During the first cycle of PCR, fragments containing flanking genomic DNA are amplified by a primer spanning the transgene/3’LTR. The longer cassette strand does not prime because its complementary shorter strand has not ligated; whereas the shorter cassette strand does not prime because only 10 nucleotides are complementary to the longer cassette strand, resulting in a high annealing temperature. This cassette design limits the amplification of non-flanking genomic DNA and reduces PCR blocking by the shorter cassette strand. Subsequent cycles are primed by both the transgene/3’LTR primer and the longer cassette strand.

### Inverse PCR

Inverse PCR was performed as previously described [15, 16]. In brief, DNA fragments were circularized in a dilute mixture of 1ng/μL DNA in T4 DNA ligase buffer with 1 cohesive end unit/μL T4 DNA ligase (New England Biolabs) for 16 hours at 16ºC. The ligation product was purified by ethanol precipitation. The first PCR reaction was 35 cycles, using a forward primer that was complementary to the distal 3’LTR and directed towards the flanking genomic DNA; and a reverse primer complementary to the junction between the distal transgene and 3’LTR (Fig.1B). The PCR mixture was as follows: 50 – 400ng circularized DNA template, forward and reverse primers (100nM each), dNTP (250nM), 1.25 Unit HotStart Taq DNA polymerase (Qiagen, Hilden, Germany), 1X PCR buffer and 1X Q-Solution (Qiagen), in a final reaction volume of 50μL. A 0.5μL aliquot of the first PCR product was amplified in a nested PCR reaction for 35 cycles, and tailing sequences were added for barcoding and Nanopore sequencing. All PCR reactions were performed on a Bio-Rad T100™ thermal cycler (Bio-Rad Laboratories, Hercules, CA). Primer sequences and thermocycling conditions are listed in Table 1.

**Table 1.**
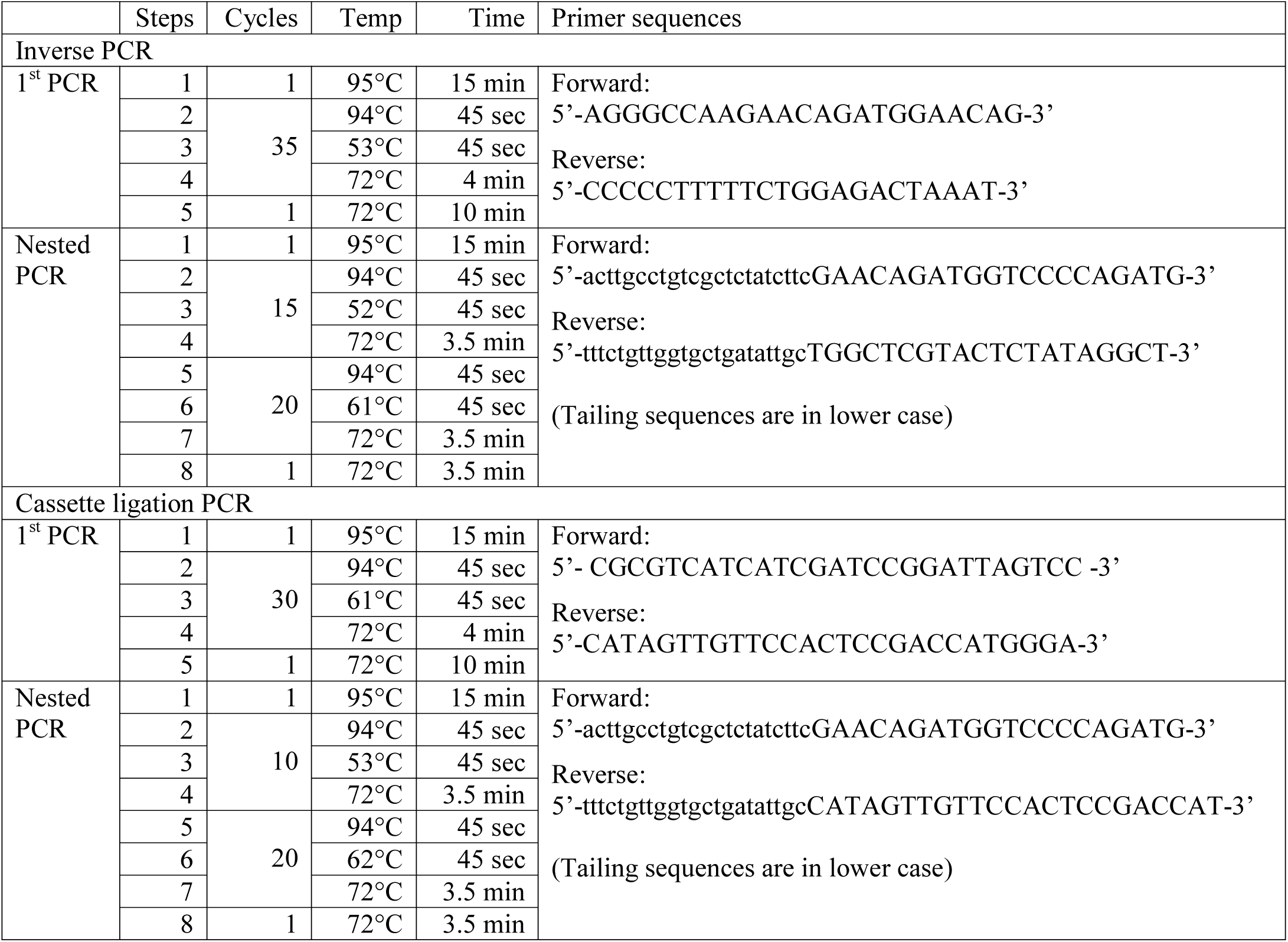
Thermocycling conditions and primer sequences

### Cassette ligation PCR

Cassette ligation PCR was performed according to the schema on Fig.1C. The restriction enzymes generated 5’ overhangs. A single dideoxy-CTP (ddCTP) was filled in to prevent elongation of the recessed 3’ ends using the following reaction mixture: *NcoI/BspHI* digested DNA 16.7ng/μL, dideoxy-CTP 33μM (GE Healthcare, Chicago, IL), DNA polymerase I, large (Klenow) fragment 0.167U/μL (New England BioLabs) in CutSmart buffer (New England BioLabs) in a final reaction volume of 15μL. The mixture was incubated at 25°C for 30 minutes, heat inactivated at 75°C for 20 minutes, and purified by ethanol precipitation.

#### Cassette ligation

Linker cassettes were made by annealing two single-stranded DNAs: 5’-ATGTCCCATGGTCA-3’ and 5’-CATAGTTGTTCCACTCCGACCATGGGA-3’ (both at 20μM) in Tris-HCl (50mM) with MgCl_2_ (5mM). The mixture was heated to 95°C for 5 minutes then gradually cooled to room temperature by turning off the heating block. The linker cassettes were aliquoted, stored at −20°C, and used without repeated freeze/thaw. The ddCTP-filled in DNA fragments were ligated to the linker cassettes using the following reaction mixture: DNA 5ng/μL, linker cassette 500nM, and T7 DNA ligase 30U/μL in T7 ligase buffer (New England BioLabs). The mixture was incubated at 25°C for 30 minutes, heat inactivated at 65°C for 10 minutes, and purified by ethanol precipitation. Because the 3’ recessed end of the DNA fragments had a ddCTP, this reaction resulted in ligation of the longer linker cassette strand and a nick in the shorter linker cassette strand. The first PCR reaction was 30 cycles in 50μL: the forward primer was complementary to the junction between the transgene and the 3’LTR to minimize non-informative reads from 5’LTR priming; and the reverse primer was identical to the longer linker cassette strand. The unligated linker cassette strand was short (14 nucleotides) relative to the ligated cassette strand (27 nucleotides), which reduced unwanted PCR priming of non-flanking genomic fragments. A 0.5μL aliquot of the first PCR product was amplified 30 cycles in a nested PCR reaction (50μL) which included tailing sequences for barcoding and Nanopore sequencing.

### Fragment size estimation by agarose gel electrophoresis

Fragment size estimation was performed by agarose gel electrophoresis and imaged on Vilber Lourmat Ebox CX5 (Vilber Lourmat). The nested PCR products (8μL) were run on 1% agarose gel. To check for specificity of the amplicons, the PCR products were digested with *SmaI*, which has cut sites within the 3’LTR and distal transgene, generating DNA fragments of 244bp and 391bp from inverse PCR amplicons and fragments of 244bp from cassette ligation PCR amplicons (Fig. 2).

**Figure 2.**
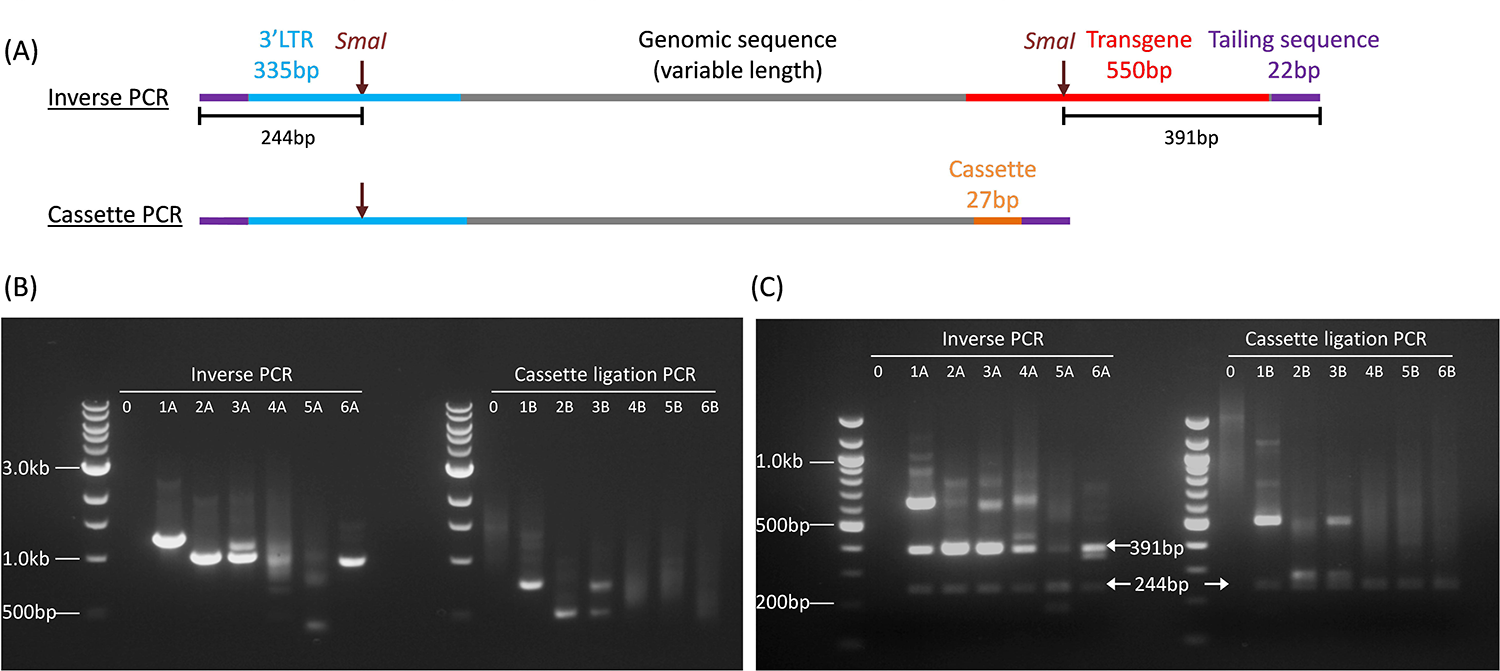
Schematic and agarose gel electrophoresis of PCR amplification products. (A) Schematic representations of PCR amplicons from inverse PCR (upper) and cassette ligation PCR (lower). The theoretical minimum lengths including the tailing sequences (22 bp × 2) are 929bp and 406bp, respectively. (B) 1% agarose gel electrophoresis of inverse PCR (left) and cassette ligation PCR (right) products. Lane 0: Non-transduced genomic DNA control. A small proportion of bands (especially Sample 5A) were smaller than the theoretical minimum lengths. (C) 2% agarose gel electrophoresis of PCR products after *SmaI* digestion. Arrows point to the expected specific bands.

### Nanopore sequencing

Sequencing libraries were prepared according to 1D PCR barcoding amplicons protocol from Oxford Nanopore Technologies (ONT, Oxford UK). Firstly, PCR amplicons from inverse PCR and cassette ligation PCR were barcoded using PCR barcodes provided by the supplier. Barcoding PCR mixture was as follow: 24μL of nested PCR product (0.5nM), 25μL of LongAmp Taq master mix (New England Biolabs) and 1μL of PCR barcode (ONT). Reactions were amplified in a thermal cycler with the following conditions: 95°C for 3 minutes, 15 cycles of amplification at 95°C for 15 seconds, 62°C for 15 seconds and 65°C for 4 minutes, followed by final extension of 65°C for 1 minute. Barcoded PCR products were purified using 0.8 X Agencourt AMPure XP beads (Beckman Coulter, Brea, CA). Concentrations of purified PCR products were measured using Qubit High Sensitivity Kit (Thermo Fisher, Waltham MA). Based on the concentrations, PCR products were pooled using 10 times more polyclonal clinical samples as compared to oligoclonal samples to increase sequencing coverage. Sequencing libraries were prepared from pooled barcoded PCR products according to manufacturer’s instruction. Briefly, end repair was performed in the following mixture: 48μL of pooled PCR product, 3.5μL of NEBNext FFPE DNA repair buffer, 2μL of NEBNext FFPE DNA repair mix, 3.5μL of Ultra II End-prep reaction buffer and 3μL of Ultra II End-prep enzyme mix (all from New England Biolabs); the mixture was incubated at 20°C for 15 minutes and 65°C for 15 minutes. End repaired products were purified with 1X Ampure XP beads (Agencourt) and eluted in 60μL of Nuclease-free water. Purified end-repaired products were ligated with 25μL Ligation Buffer (ONT), 10μL NEBNext Quick T4 DNA Ligase (New England Biolabs) and 5μL Adapter Mix (ONT) and incubated at room temperature for 30 minutes. Ligated products were purified with 0.4X Ampure XP beads (Agencourt) and Short Fragment Buffer (ONT) and final sequencing library was eluted in Elution Buffer (ONT). Sequencing library was loaded into a single ONT PromethION flowcell and sequenced for 40 hours.

### Sequence Data Analysis

A flow diagram for sequence data analysis is outlined in Fig. 3. Analysis parameters and custom scripts are provided in **Supplementary information**. Sequencing reads were base-called and demultiplexed using Guppy basecaller version 2.3.7 (ONT). Reads were classified based on the read quality score as pass (≥7) or fail (< 7) by the basecaller. Adapters and sample barcodes were trimmed using Porechop version 0.2.4 (https://github.com/rrwick/Porechop). Reads that were shorter than the combined lengths of the predicted flanking sequences of the amplicons (885bp for inverse PCR and 362bp for cassette ligation PCR) were excluded from analysis. For inverse PCR, filtered reads were aligned to hg38 genome and both flanking sequences (i.e. 3’LTR and distal transgene sequence), with masking of the distal transgene sequence using BWA mem (version 0.7.15) [18]. For cassette ligation PCR, the flanking cassette sequence was short and difficult to align; hence this was first trimmed with Porechop and the trimmed reads were then aligned to hg38 genome and flanking 3’LTR with BWA mem (version 0.7.15) [18]. Note that as a result of cassette trimming, the final reads for cassette ligation PCR were 27bp shorter than the original filtered reads. To determine which of the two ends represented the 3’LTR-genome junction, we developed a tool called Flankdetect (https://github.com/mdcao/japsa) (version 1.9-10b) (**Supplementary information**), which identifies reads which contain the flanking sequences and reports the integration site that flanks the junctional 3’LTR sequence. This tool also assigns clonality by clustering reads based on the flanking integration sites. Reads which have integration sites within 10bp of each other are clustered together. During clustering, alignments that were not primary alignment and less than mapping quality of 20 were excluded from further analysis. To eliminate false clusters, Bedtools (version 2.26.0) merge option was used to merge any overlapping clusters which contain reads that align to the same region. The integration site that is most frequently observed within all the reads that are merged together is retained as the integration site of the clone.

**Figure 3.**
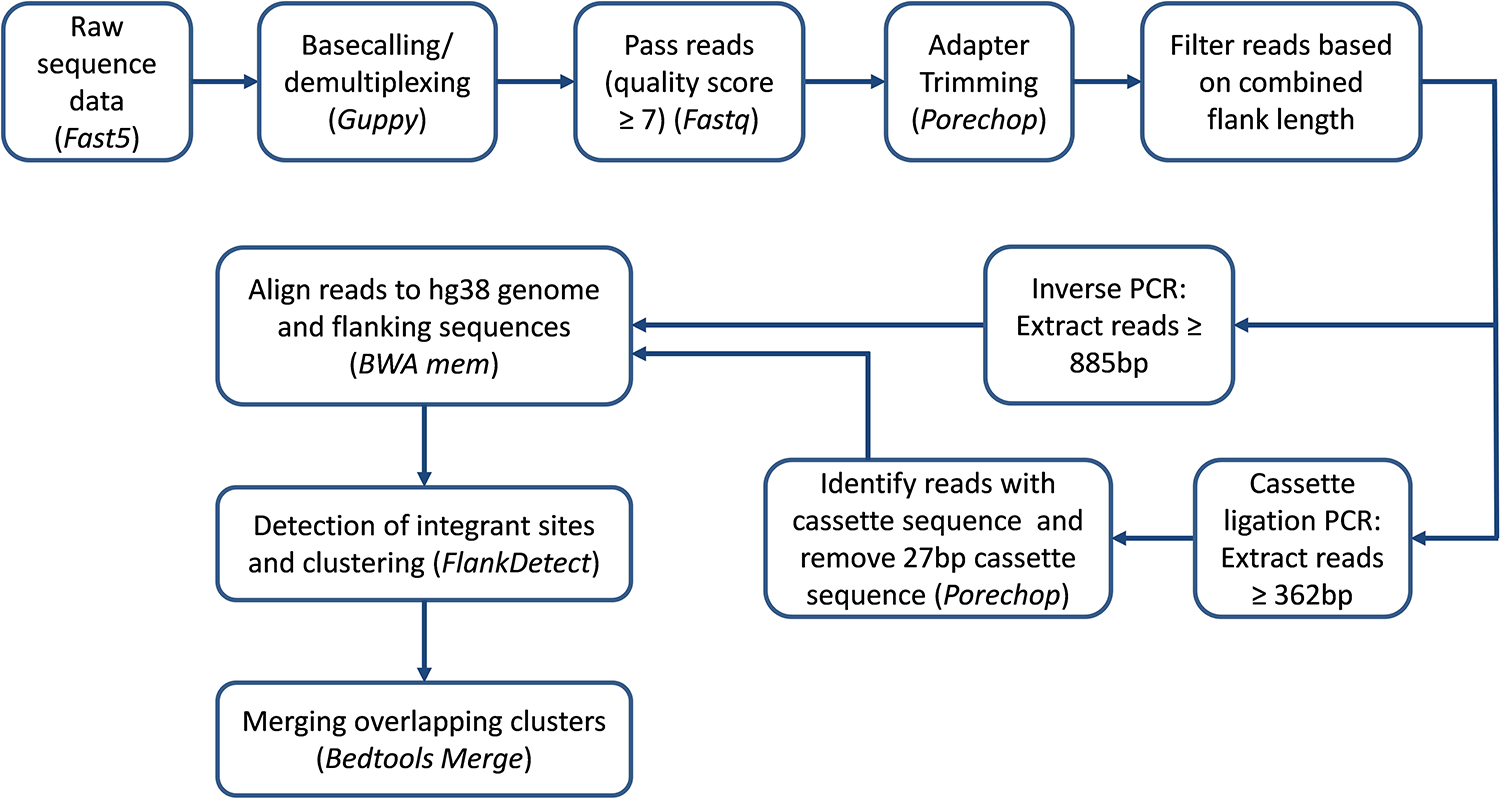
Flowchart for Nanopore sequencing analysis. Summary of sequencing data analysis. Tools used for analysis are italicized.

### Annotation of Integration Site

Genomic locations of transcription start site (TSS), exonic regions, intronic regions and intergenic regions were extracted from GENCODE version 28 gene annotation file using custom scripts (**Supplementary information**). Genomic annotation of integration site was performed using Bedtools (version 2.26.0) intersect option. Genomic distance of the integration site from TSS was calculated using custom scripts (**Supplementary information**).

## Results

### Amplification of flanking sequence DNA by inverse PCR and cassette ligation PCR

The amplicons from inverse PCR and cassette ligation PCR were predicted to have a sandwich structure: the flanking genomic sequence in the middle, which would be contiguous with the 3’LTR on one end, and the sticky-end ligated distal transgene sequence or linker cassette sequence on the other end (Fig.2A). The theoretical minimum lengths of productive amplicons, including the tailing sequences, were 929bp for inverse PCR and 406bp for cassette ligation PCR. We performed flanking sequence amplification on SFG.iCasp9.2A.ΔCD19-transduced Jurkat cell clones and polyclonal clinical samples from our previously published phase I clinical trial [15]. As expected, inverse PCR and cassette ligation PCR on transgenic Jurkat cell clones (clones 1 and 2) resulted in single PCR fragments (Fig.2B: lanes 1A, 1B, 2A, 2B); and a 1:1 mixture of the two Jurkat cell clones resulted in a combination of the two bands (lanes 3A and 3B). Amplification of a polyclonal SFG.iCasp9.2A.ΔCD19-transduced clinical cell product yielded a polyclonal smear (lanes 4A and 4B), as did amplification of patient PBMC collected at Day 369 after T cell infusion (lanes 5A and 5B). However, amplification of patient PBMC from Day 1332 after T cell infusion showed a dominant band with inverse PCR and a polyclonal smear with cassette ligation PCR (lanes 6A and 6B), suggesting a possibility of bias in DNA fragment circularization or PCR amplification with the former. The inverse PCR amplicons in lanes 4A and 5A also contained bands that were shorter than the minimum theoretical length of 929bp. These were later shown on Nanopore sequencing to contain 3’LTR and transgene sequences without any intervening genomic DNA, which was consistent with the circularization of degraded DNA fragments near the 3’LTR.

### Specificity of PCR amplification

A combination of two 6-cutters (*NcoI* and *BspHI*) is expected to cut at approximately every 2,048 bp (i.e. 4^6^/2), generating an estimated 1.5 × 10^6^ DNA fragments per diploid human cell, which has a genome of approximately 3 ×10^9^ bp. Given that a majority of transgenic T cells carry only a small number of transgene inserts per cell, this would mean that only a tiny fraction (<0.005%) of the DNA fragments would include a transgene flanking sequence. Non-specific amplification of non-flanking sequences is therefore a challenge. As shown in Fig.2A, specific PCR amplicons would contain the predicted flanking sequences, which contained *SmaI* restriction sites within the 3’LTR flanking sequence following both inverse PCR and cassette ligation PCR, and an additional *SmaI* restriction site at the transgene end for inverse PCR. Thus, *SmaI* restriction enzyme digestion of specific PCR amplicons is anticipated to yield a 244bp DNA fragment from the 3’LTR end and, in the case of inverse PCR, an additional 391bp DNA fragment from the transgene end, along with flanking genomic DNA fragments of variable lengths. We digested the PCR products with *SmaI* and confirmed that the amplicons were specific, with the generation of 244bp and 391bp fragments from inverse PCR and a 244bp fragment from cassette ligation PCR (Fig.2C).

### Nanopore output and read filtering

All 12 paired PCR products from the 6 samples were pooled into one Nanopore sequencing run. In order to ensure adequate read-depth, the amount of input DNA was limited to the equivalent of less than 10,000 transduced cells per sample (see Table 2). Because the Jurkat samples were clonal and hence required fewer total reads, they were pooled with the polyclonal patient samples at a ratio of 1:10. The sequencing run produced 23.4 million reads in total and 59% of these reads were classified as passing quality score (read quality ≥ 7). Approximately 6% of the pass reads could not be demultiplexed and were excluded. Most of the samples had > 90% of reads that were longer than the minimum theoretical length of 885bp for inverse PCR and 362bp for cassette ligation PCR after adapter trimming. Table 3 summarizes the number of reads retained after each data analysis step.

**Table 2.**
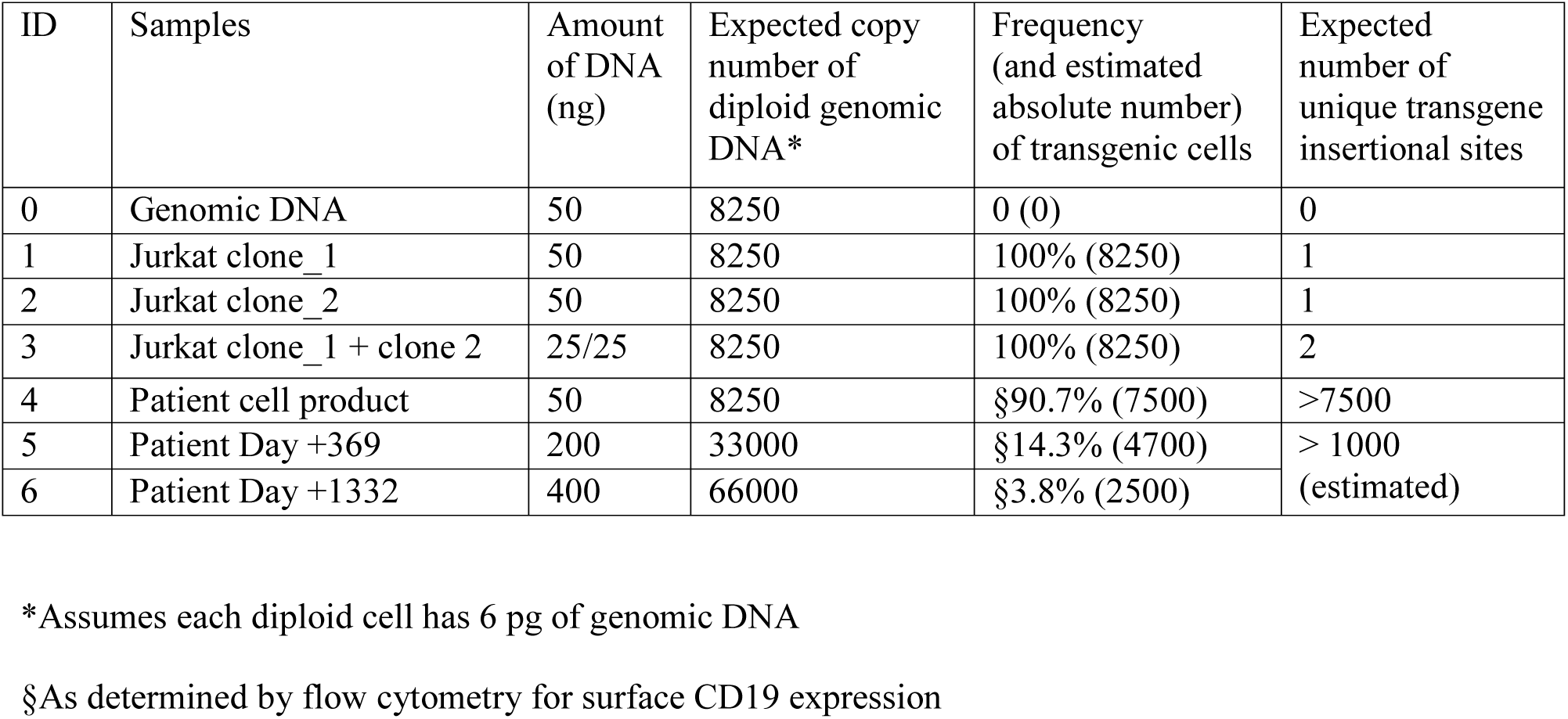
Samples for Nanopore sequencing

**Table 3.** Number and proportion of reads after filtering and alignment

|  |  | Sample ID | Pass reads | Adapter Trimmed Reads |  | Reads > Minimum theoretical length |  | Reads with both flanks |  | Aligned reads |  |
| --- | --- | --- | --- | --- | --- | --- | --- | --- | --- | --- | --- |
|  |  |  | Number of reads | Number of reads | % of pass reads | Number of reads | % of adapter trimmed reads | Number of reads | % of reads > minimum theoretical length | Number of reads | % of reads with both flanks |
| Inverse PCR | Jurkat cells | 1A | 56,043 | 50,396 | 89.9% | 48,451 | 96.1% | 44,776 | 92.4% | 44,475 | 99.3% |
|  |  | 2A | 272,138 | 249,196 | 91.6% | 237,487 | 95.3% | 232,040 | 97.7% | 218,894 | 94.3% |
|  |  | 3A | 157,759 | 146,228 | 92.7% | 139,974 | 95.7% | 136,263 | 97.3% | 129,181 | 94.8% |
|  | Clinical samples | 4A | 1,949,817 | 1,800,452 | 92.3% | 581,723 | 32.3% | 519,444 | 89.3% | 327,685 | 63.1% |
|  |  | 5A | 3,102,548 | 2,889,377 | 93.1% | 231,456 | 8.0% | 213,196 | 92.1% | 146,956 | 68.9% |
|  |  | 6A | 2,868,949 | 2,645,501 | 92.2% | 2,377,721 | 89.9% | 2,338,293 | 98.3% | 1,724,250 | 73.7% |
| Cassette Ligation PCR | Jurkat cells | 1B | 91,891 | 84,230 | 91.7% | 78,796 | 93.5% | 73,476 | 93.2% | 70,763 | 96.3% |
|  |  | 2B | 150,027 | 135,417 | 90.3% | 129,610 | 95.7% | 125,634 | 96.9% | 118,086 | 94.0% |
|  |  | 3B | 131,303 | 111,739 | 85.1% | 110,416 | 98.8% | 102,695 | 93.0% | 97,443 | 94.9% |
|  | Clinical samples | 4B | 1,488,028 | 1,373,853 | 92.3% | 1,272,524 | 92.6% | 1,234,596 | 97.0% | 856,189 | 69.3% |
|  |  | 5B | 1,333,591 | 1,203,536 | 90.2% | 1,072,963 | 89.2% | 1,040,570 | 97.0% | 726,298 | 69.8% |
|  |  | 6B | 1,228,503 | 1,137,012 | 92.6% | 945,077 | 83.1% | 926,182 | 98.0% | 435,600 | 47.0% |

### Sequence alignment and clustering

Greater than 94% of reads containing the expected flanking sequences from the transduced Jurkat cell lines (Samples 1, 2 and 3) could be uniquely aligned to the genome (Table 3). The genome alignment rate was lower for the polyclonal clinical samples, with 47% to 74% (median 69%) of reads with the expected flanking sequences being successfully aligned. A majority of reads that could not be aligned to genomic DNA were only marginally longer than the minimum theoretical lengths of the PCR amplicons and were therefore too short to be uniquely aligned (Fig.4A). Long unaligned reads generally lacked the anticipated flanking 3’LTR/transgene or cassette sequences and therefore represented non-specific amplicons. A significant proportion of inverse PCR amplicons in Samples 4 and 5 were below the minimum theoretical lengths. Alignment of 3 dominant clones (560 bp, 388 bp and 318 bp) showed that these consisted of the terminal regions of the flanking 3’LTR and transgene sequences, without any intervening genomic DNA (Fig.4B), in an orientation that was consistent with the circularization and PCR amplification of degraded DNA fragments within the transgene / 3’LTR region (Fig.4C).

Aligned reads with 3’LTR-genome junctions that were within 10 bp of each other were clustered together using our customized tool, Flankdetect. The 4,895,820 reads were grouped into 12,186 clusters. Each cluster consisted of 1 to >10^6^ reads: 59% of clusters had single reads, 24% of clusters had 2 – 10 reads, 7% of clusters had 11 – 100 reads, and the remaining 10% of clusters had >100 reads (Fig.4D). For clusters with 2 to 4 reads, the 3’LTR -genome junction was identical in 24% of the clusters. The junction varied by 1 nucleotide in 28% of clusters, and by 2 – 3 nucleotides in 18% of clusters (Fig.4E). Overall, for clusters with 2 to 4 reads, the spread of 3’LTR -genome junctions was within 5 nucleotides in 88% of the clusters, but a few clusters had a spread of 10 to 15 nucleotides. For clusters with ≥ 5 reads, we defined R80 as the span of 3’LTR -genome junction that contained at least 80% of reads within the cluster: 15% of clusters had R80 of 0 nucleotides, meaning 80% of the reads had identical 3’LTR-genome junction; and 80% of clusters had R80 of 1 nucleotide (Fig. 4F). Overall, for clusters with 5 or more reads, 97% had an R80 of ≤5 nucleotide, meaning 80% of reads within the clusters were within a span of 5 nucleotides, although there were isolated clusters with a spread of up to 40 nucleotides.

**Figure 4.**
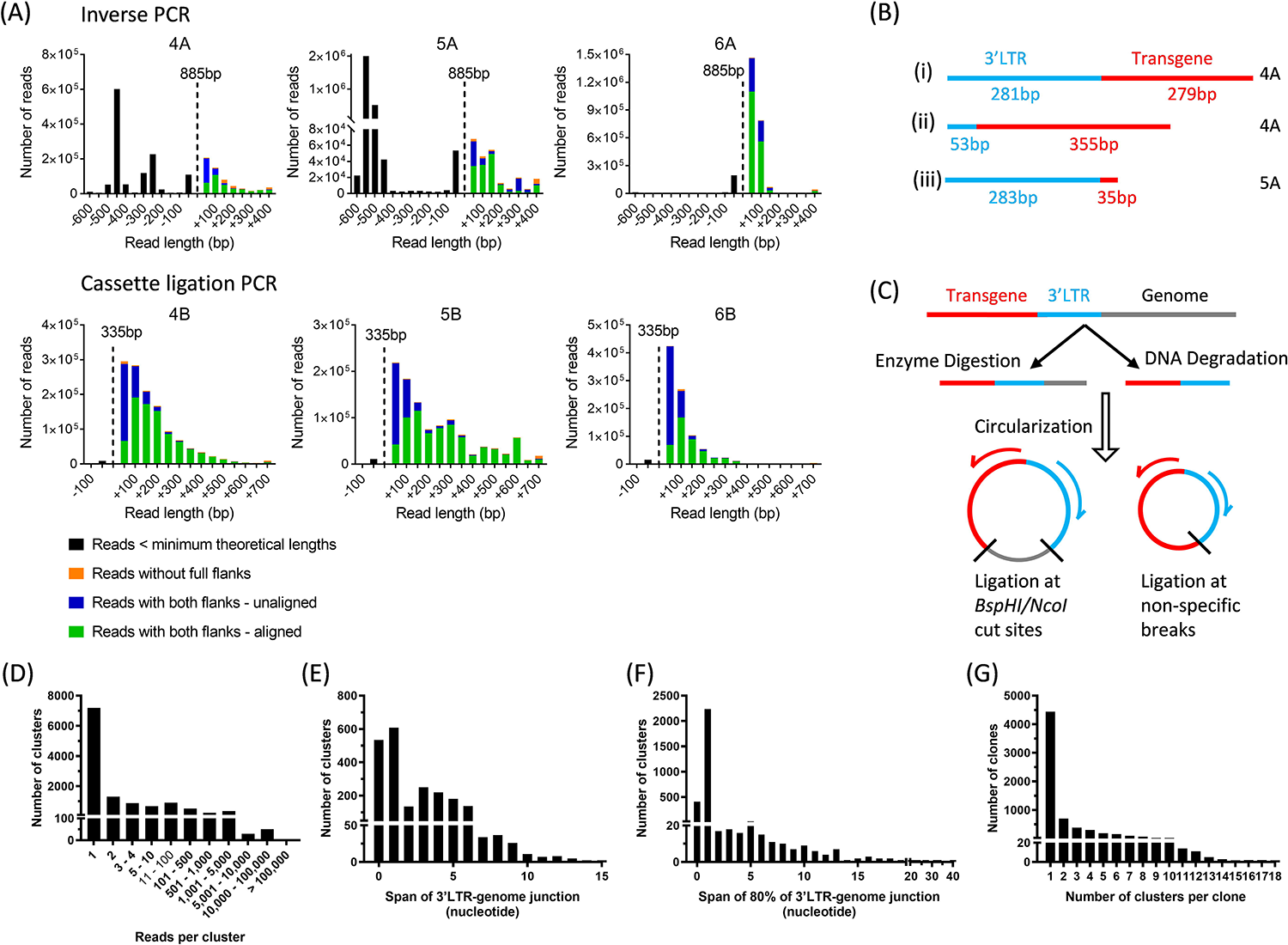
Read alignment and clustering. **(A)** Read-length distribution for the polyclonal clinical samples by inverse PCR (top row) and cassette ligation PCR (bottom row) relative to their respective minimum theoretical lengths of 885 bp and 335 bp (after cassette trimming). Shown are reads that were below the minimum theoretical length (black), reads that were above the minimum theoretical length but did not have the expected flanking sequences (orange), and reads with the expected flanking sequences and were aligned (green) or unaligned (blue) to genomic DNA. (B) Schematic representation of the dominant short inverse PCR amplicons in Sample 4A (i and ii), and 5A (iii). Note that the junctions between the 3’LTR and transgene were formed by circularization of DNA fragments and were not the native junction. (C) Schematic representation of the formation of short amplicons from DNA fragments. (D) Distribution of number of reads per cluster. Shown are data for all 12 samples. (E) Span of 3’LTR-genome junction for clusters that contained 2 to 4 reads. ‘0’ indicates identical 3’LTR-genome junction for all reads within the cluster. (F) Span of 3’LTR-genome junction that includes 80% of reads in clusters that include 5 or more reads. (G) Number of clusters per clone after merging clusters with overlapping read alignment.

We observed that a proportion of the clusters were very proximate to each other. In some cases, the read alignments were very closely matched but the clusters were considered separate because the 3’LTR-genome junction was assigned to opposite ends of the read, which would suggest an underlying alignment error. In order to avoid splitting clonal integration sites into multiple artificial clones, clusters with overlapping read alignments were merged and considered as belonging to the same clone. Using this merging algorithm, we identified 6,410 unique vector integration clones: 4,440 clones (69%) consisted of only 1 cluster, 697 clones (11%) consisted of 2 clusters, 383 clones (6%) consisted of 4 clusters, and the remaining 890 clones (14%) consisted of 4 – 18 clusters (Fig.4G).

### Clonal composition by inverse PCR and cassette ligation PCR

Samples 1 and 2 each contained a unique vector integration site, which were detected by both inverse PCR and cassette ligation PCR. Note that both samples had a degree of subclonal contamination, which was more prominent on cassette ligation PCR than inverse PCR. Sample 3 consisted of an equal mixture of Samples 1 and 2 but there was a dominance of the integration site from Sample 2, with the bias being more pronounced by inverse PCR (Fig.5A). The clinical samples were polyclonal: the cell product (Sample 4) was anticipated to be the most clonally diverse, with some degree of clonal dominance emerging in the post-infusion patient samples (Samples 5 and 6) [15]. The number of unique vector integration sites identified in Samples 4, 5 and 6 were 979, 196 and 602 by inverse PCR and 2258, 1251 and 669 by cassette ligation PCR, with the starting materials containing approximately 7500, 4700 and 2500 *iCasp9*-transduced cells. In all three samples, clonal skewing was more pronounced with inverse PCR than cassette ligation PCR: the top 10 integration sites in Samples 4, 5, and 6 accounted for 49%, 93% and >99%, respectively, of the total aligned reads by inverse PCR; as compared to 7%, 58% and 76% of the total aligned reads by cassette ligation PCR (Fig.5B). A proportion of the integration sites were detected by both techniques but their percentage representation within the sample differed between the two techniques, and a majority of integration sites were mapped by only one of the techniques (Fig.5C). The very large clone representing 97% of reads in Sample 6 by inverse PCR, represented only 0.1% of the reads by cassette ligation PCR. This clone also had higher clonal representation by inverse PCR relative to cassette ligation PCR in Samples 4 and 5 (0.08% versus 0.02%; and 0.44% versus 0.15%), suggesting amplification bias.

**Figure 5.**
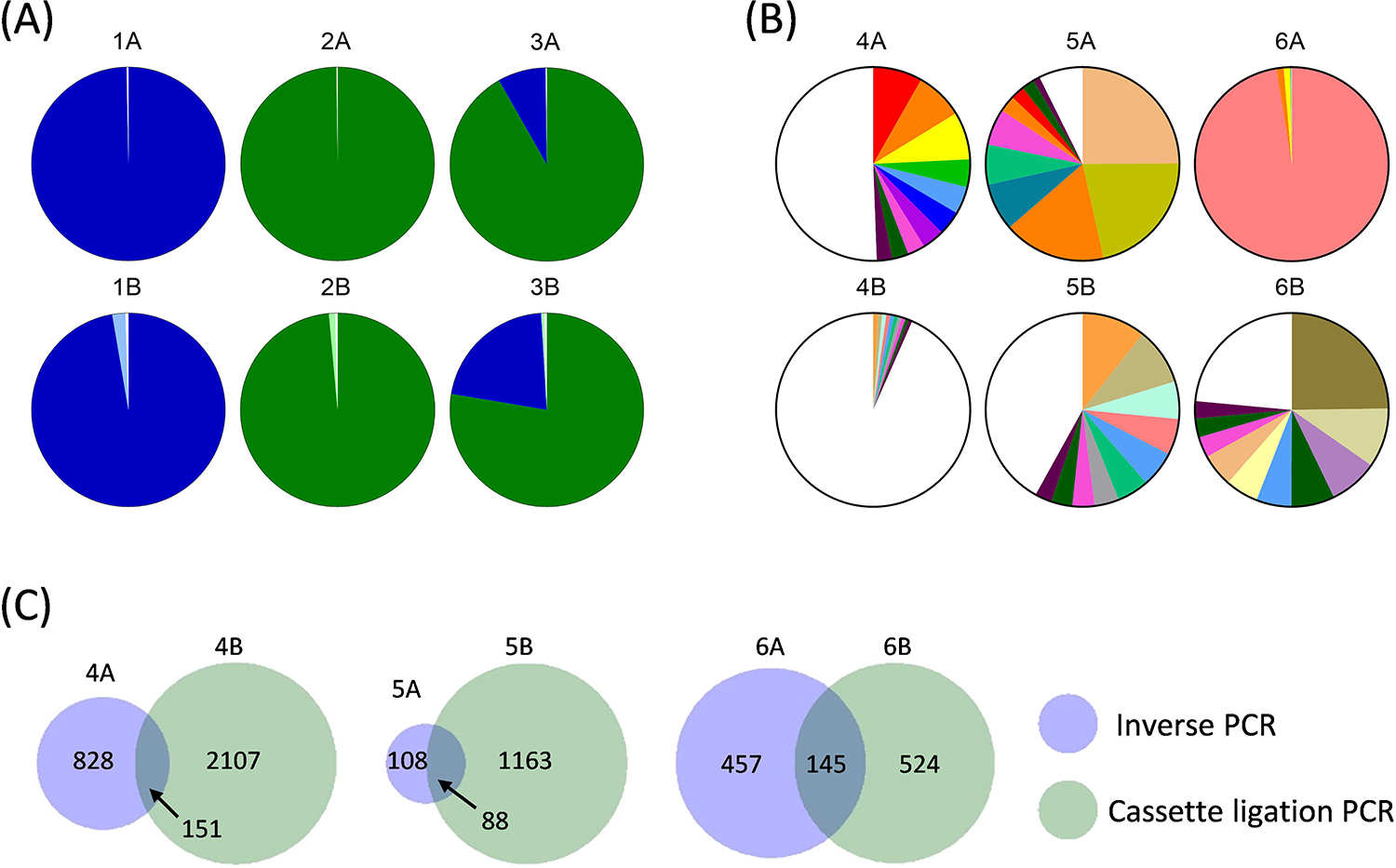
Clonal composition of vector integration sites. (A) Distribution of vector integration sites as a proportion of total aligned reads in *iCasp9*-transduced Jurkat cell clones. Shown are paired analysis by inverse PCR (top row) and cassette ligation PCR (bottom row). Samples 1 and 2 were two separate Jurkat cell clones, and Sample 3 was a 1:1 mix of Samples 1 and 2. Green and blue indicate different clones. Light green and light blue indicate the subclones. (B) Distribution of vector integration sites as a proportion of total aligned reads in polyclonal clinical samples. Shown are paired analysis by inverse PCR (top row) and cassette ligation PCR (bottom row). Sample 4 was *iCasp9*-transduced cell product; and Samples 5 and 6 were PBMC obtained at day +369 and +1332 after cell infusion, respectively. The top 10 clones for each sample are shown in color; with the remainder represented in white. The color representation is random and does not correspond to the same clones across different pie charts. (C) Overlap of unique vector integration sites identified by inverse PCR and cassette ligation PCR in the polyclonal clinical samples. Figure 5A and B ere plotted with Prism 7, GraphPad, and Figure 5C was plotted with R version 3.5.3 (venneuler package).

### Location of integration sites within the genome

Vector integration sites were identified within all chromosomes, with higher representations within the larger chromosomes, as these represented a larger component of the total genome (Fig.6A). A majority (61 – 68%) of integration sites were intragenic, with 55 – 58% within the introns and 6 – 12% within the exons; with the remaining 32 – 39% of integration sites being intergenic (Fig.6B). There was a predilection for vector integration near transcription start sites (TSS) (Fig.6C), which was consistent with other reports [3, 5, 6].

**Figure 6.**
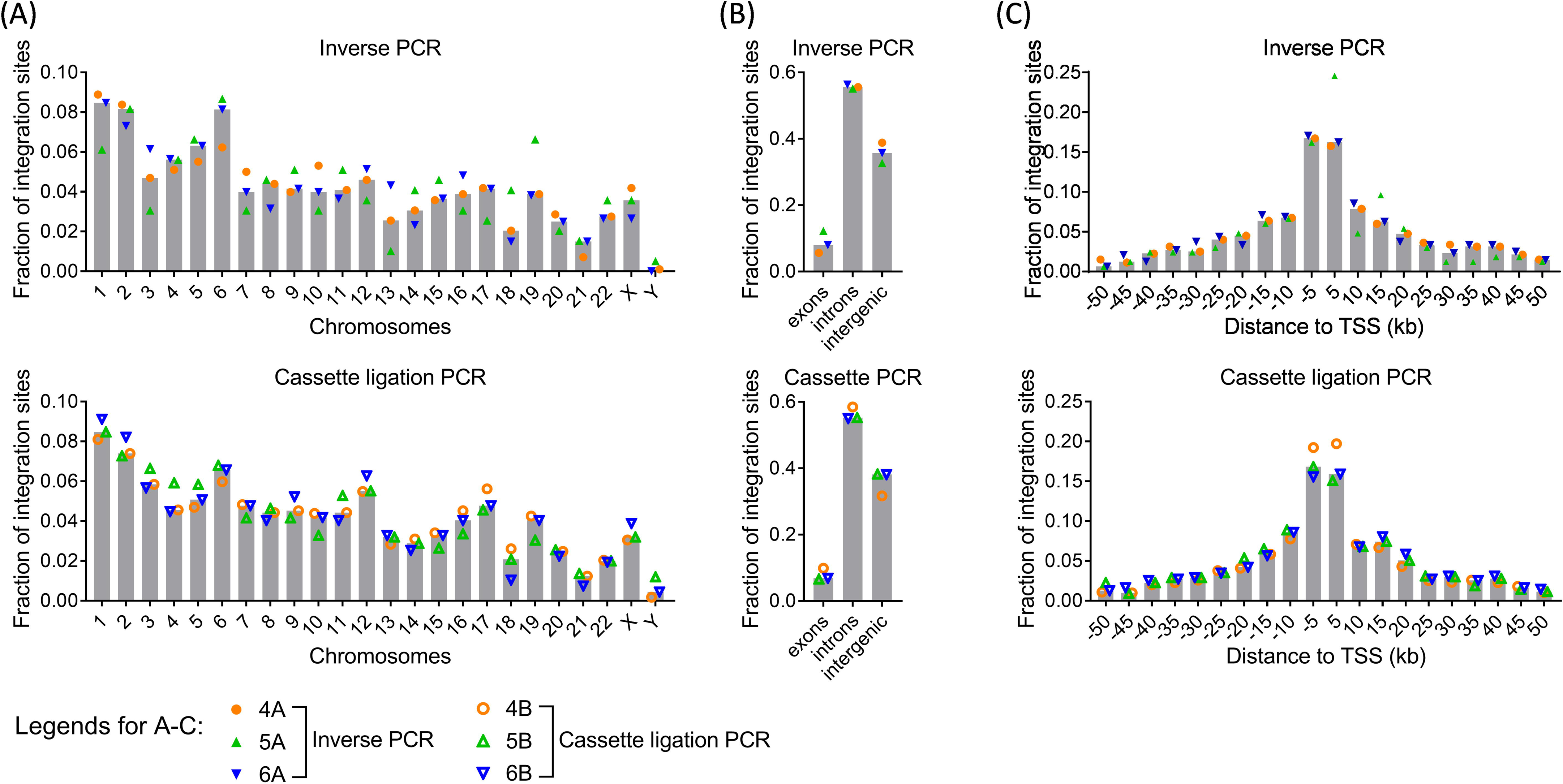
Annotation of vector integration sites. Distribution of unique vector integration sites within the genome as detected by inverse PCR (top) and cassette ligation PCR (bottom). Integration sites are annotated by: (A) chromosomes, (B) gene coding regions (exons, introns and intergenic), and (C) distance to transcription start sites (TSS). Median values were presented (Prism 7, GraphPad).

## Discussion

Analysis of vector integration sites is currently restricted to highly specialized academic institutions where it is used largely as a research tool. However, gene-modified cellular therapeutics, especially CAR T cells, are rapidly entering routine clinical practice and vector integration site analysis would be required from time-to-time to monitor for the emergence of dominant clones. Many smaller clinical centers do not have access to expensive sequencing instrument and, furthermore, whilst short-read next generation sequencing is cost effective for large batches, the cost per batch is very high. As a result, centers with low sample volumes will need to either run smaller batches at significantly higher cost per sample, or accumulate samples over a very long period of time for cost-effective batching, which reduces the timeliness of the analysis. In this paper, we showed that Nanopore sequencing platform could be successfully utilized to map polyclonal vector integration sites.

We used two different methods to amplify flanking genomic DNA: inverse PCR and cassette ligation PCR. Inverse PCR was slightly less demanding technically but has a higher potential for bias because efficient circularization is limited to DNA fragments of 300bp to 3kb, and secondary structure formation can further limit circularization efficiency. There is also competition between intramolecular circularization and intermolecular ligation, although this can be partially minimized using a dilute DNA ligation mix. In order to reduce bias, we adapted a previously described cassette ligation PCR method [19] to amplify the flanking genomic sequences with high efficiency and specificity. This method had significantly less bias compared to the inverse PCR method but it was not completely bias-free as an equal mix of the two Jurkat clones did not produce equal representations of reads from each clone, although was less skewed than inverse PCR.

Our analysis considered clusters with any degree of overlap in read alignments as belonging to the same vector integration site. It is possible that this could erroneously merge proximate but distinct vector integration sites, resulting in an underestimation of clonal diversity. We have developed our clustering protocol to err on the side of over-clustering because a key function of vector integration site analysis is to detect clonal dominance, which can sometimes indicate autonomous growth or insertional mutagenesis, and it is therefore important to avoid splitting single integration sites into multiple artificial clones. The number of unique integration sites mapped in our study was consistent with those using LAM-PCR followed by next generation short-read sequencing, which typically detects around 200 to 8,000 unique integration sites from 10^3^ to 10^6^ transduced T cells [3, 5, 6]. However, in our analysis of polyclonal clinical samples using inverse PCR and cassette ligation PCR in parallel, only 7% of unique vector integration sites could be detected by both methods. This low level of overlap was likely a result of both PCR amplification bias and sampling artefact. Gene-modified T cells, including *iCasp9*-transduced T cells, have been shown to be highly polyclonal by T cell receptor [15, 20] and vector integration site analysis [3, 15]. We and others have detected 10^3^ to 10^4^ unique clonotypes within aliquots of infused cell products and post-infusion patient samples in T cell therapy trials using gammaretroviral vectors [3, 6, 15]. The actual clonal richness is likely much higher if a very large number of cells can be sampled and sequenced at depth without bias. The likelihood of a particular clone being randomly sampled twice is a function of its clonal frequency and the sample size. In this study, the amount of input DNA in the polyclonal clinical samples represented < 0.001% of the total pool of transduced cells; hence, our observation that only a small proportion of clones could be detected by both inverse PCR and cassette ligation PCR was consistent with the anticipated low likelihood of a particular clone being randomly sampled into both reaction mixtures. Nonetheless, the large differences in the clonal size of the identified overlapping clones would suggest that there was also significant contribution from amplification or sequencing bias.

In studies using gene-modified T cells, clonality can also be assessed by T cell receptor clonotype analysis. Although this remains an expensive assay, it is somewhat more accessible and can be outsourced to commercial entities. T cell receptor clonotype can provide a good indication of the clonal diversity of the transduced cells because clones bearing the same vector integration sites will bear the same T cell receptor clonotype. However, T cell receptor analysis does not provide any information on the nature of the vector integration sites and cells with the same T cell receptor clonotype can sometimes carry different vector integration sites through separate transduction events [15, 21].

The amount of data that can be generated and the feasibility to scale as required make Nanopore sequencing an attractive option for analysis of vector integration sites. In this report, we re-used an ONT PromethION flow cell and obtained 15 Gb of sequencing data, which was sufficient to analyze 6 polyclonal and 6 oligoclonal samples, with >5,500 unique integration sites identified. Based on this estimate, the smaller ONT MinION flow cell, which costs around USD 900 and capable of generating 20 – 30 Gb of sequencing data, could be used to sequence 12 polyclonal samples; whereas the larger ONT PromethION flow cell, which costs around USD 2,000 and capable of generating around 80 Gb of sequencing data, would be suitable for larger batches. There is also now a very small ONT Flongle that is capable of 1Gb of sequencing data at a cost of USD 160 each, which may be suitable for single sample analysis. Another distinct advantage of Nanopore sequencing is its low cost of entry as it does not require any dedicated sequencing instrument, making it highly feasible for smaller centers. These features are of immense relevance in the context of current developments in the CAR T cell field, which have seen their emergence from large dedicated academic research centers into routine implementation in smaller clinical centers where the ability to perform vector integration site analysis in a timely manner, with flexibility of scale, can be clinically important. The total assay time is under four days: one-and-a-half days to complete ligation and nested PCR, and two days for library preparation, sequencing and data analysis.

The main drawback of Nanopore sequencing is its high error rate. This limits its ability to confidently map the 3’LTR-genome junction at single nucleotide resolution. Nonetheless, a majority of clusters could be resolved within a 5-nucleotide span and 94% of clusters with 5 or more reads could be resolved within 1 nucleotide span. The resolution will likely improve with evolution in the technology platform and refinement in the assay design and data analysis. At present, the higher error rate also meant that relatively long stretches of flanking DNA sequence are required for successful alignment. The length of the flanking DNA sequence is pre-determined by the position of the restriction site and hence integration sites that are very close to the restriction sites will always be very difficult to align. However, it may be possible to perform parallel analysis using different sets of restriction enzymes or tagmentation without restriction enzymes to increase the proportion of aligned reads [5, 6].

In summary, we have developed a readily accessible, highly scalable, low cost and low capital method to analyze vector integration sites within a polyclonal sample using Nanopore sequencing. This platform has the potential to become a practical alternative to short-read next generation sequencing, especially for smaller clinical centers with low volume throughput where flexibility of scale and timeliness of results are important.

## Declarations

This trial was prospectively approved by the Human Research Ethics Committee (Institutional Review Board) of Royal Brisbane and Women’s Hospital. It was conducted in accordance with the Declaration of Helsinki and the Australian National Statement on Ethical Conduct in Human Research. Written informed consent was obtained from all participants. The trial was prospectively registered at www.anzctr.org.au as ACTRN12614000290695.

## Acknowledgements

This work was supported by a Project Grant (APP1053135) from the National Health and Medical Research Council (NH&MRC, Australia) and Royal Brisbane and Women’s Hospital Foundation. SKT was supported by an NH&MRC Early Career Fellowship (APP1054786) and Metro North Hospital and Health Service Clinician Research Fellowship. LJC was supported by an NH&MRC Career Development Fellowship (APP1130084). The authors acknowledge the advice from Dr Edmund Chang, previously at the Center for Cell and Gene Therapy, Baylor College of Medicine, Houston, Texas, USA, who suggested using Nanopore sequencing for this purpose. The authors are grateful to the patients and their families for their participation in the associated phase I clinical trial.

## Author Contributions

LJC and SKT designed the study. PZ, DG and RA performed experiments. DG, SHN and LJC performed analysis of sequencing data.

## Disclosure of Interest

The authors declare no conflict of interest.

