## Supplementary Information for "Analysis of polyclonal vector integration sites using Nanopore sequencing as a scalable, cost-effective platform"

Analysis parameters and custom scripts used for sequencing analysis are provided below.

1. Basecalling and demultiplexing Nanopore sequence data with Guppy (version 2.3.7)

```
guppy_basecaller -i <input_fast5_dir> -s <output_basecalled_dir> -c dna_r9.4.1_450bps_flipflop.cfg
--qscore_filtering -q 0 --recursive --device <cuda device name>
```

```
guppy_barcoder -i <basecalled_dir> -s <output_demultiplex_dir> --barcode_kits EXP-PBC001 –r
```

2. Trimming adapters and barcodes with Porechop (version 0.2.4)

```
porechop-runner.py --discard_middle -i <input.fastq> -o <output_trimmed.fastq>
```

3. Filter reads based on read length and trimming cassette sequence
   For inverse PCR amplicons reads greater than or equal to 885bp were selected; for cassette PCR amplicons reads greater than or equal to 362bp were selected.

```
awk 'BEGIN {OFS = "\n"} {header = $0 ; getline seq ; getline qheader ; getline qseq ; if (length(seq) >=
885) {print header, seq, qheader, qseq}}' input_trimmed.fastq > length_filtered.fastq
```

   For cassette PCR amplicons, cassette sequence were trimmed from the reads using Porechop (version 0.2.4) .

```
porechop-runner.py --discard_middle -i < length_filtered.fastq> -o <output_trimmed.fastq>
```

4. Aligning reads to flank sequences and hg38 genome using BWA-MEM (version 0.7.15)

```
bwa mem -t 8 -k11 -W20 -r10 -A1 -B1 -O1 -E1 -L0 -Y -a hg38_and_flank_sequence.fasta
input_length_filtered.fastq > hg38_with_flank_align.sam
```

   NOTE:- Transgene sequence (i.e CD19 sequence) masked reference sequence was used for inverse PCR amplicon alignments.

5. Identifying flanking integration site from aligned reads
   Flank detect analysis script can be accessed from https://github.com/mdcao/japsa (version 1.9-10b) and can be deployed using script name jsa.np.flankDetect.

```
jsa.np.flankDetect -f flank_sequence.fasta –b hg38_with_flank_align.sam -r hg38.fasta -q 20 -d 10
>integration_site_clustering.txt 2>flank_log.txt
```

NOTE: q – mapping quality; d – distance of integration sites to be clustered together; integration_site_clustering.txt > reports the integration site for each read and reports the cluster which the read belongs to; flank_log.txt > reports the number of reads with either or both or none of the flanks.

6. Merging overlapping clusters based on alignment position using Bedtools (version 2.26.0)
Custom scripts were used to re-format the output from step 5 prior to merging the overlapping clusters.

```
awk 'BEGIN{FS=OFS="\t"}$6!="NA"{if (NR!=1) print $2,$3,$4,$5,$1,$6,$7,$8,$9,$10,$11,$12,$13}'
integration_site_clustering.txt | bedtools sort > integration_site_clustering_sorted.txt

bedtools merge -c 7,8,6,6 -o distinct,count,mode,stdev -i integration_site_clustering_sorted.txt >
integration_site_clustering_merge.txt
```

7. Annotation of genomic locations of vector integration site

i. Generating bed files for genomic regions

```
bedtools complement -i gencode.v28.annotation_sorted.gff -g chr_sizes_hg38.txt >
gencode.v28.annotation_intergenic.bed

awk '{if ($3 == "exon") print $1, $4-1, $5}' gencode.v28.annotation_sorted.gff >
gencode.v28.annotation_exon.bed

bedtools complement -i <(cat gencode.v28.annotation_intergenic.bed
gencode.v28.annotation_exon.bed | sort -k1,1 -k2,2n) –g chr_sizes_hg38.txt >
gencode.v28.annotation_intron.bed

awk '{if($3 == "gene") {if($7 == "-") {$4 = $5} else {$5 = $4} print}}'
gencode.v28.annotation_sorted.gtf | awk '{print $1, $4, $5, $7, $10, $14}'  >
gencode.v28.annotation_gene_TSS.bed
```

ii. Annotating integrant sites for exonic, intronic and intergenic regions
Annotation of integrant site was performed on the re-formatted output from step 6.

```
awk 'BEGIN{FS=OFS="\t"}{print $1,$6,$6+1,$2,$3,$4,$5}' integration_site_clustering_merge.txt >
integration_site_annotation.txt
```

Annotating to genomic regions

```
bedtools intersect -u -a integration_site_annotation.txt -b gencode.v28.annotation_exon.bed >
SampleName_exon.bed
```

```
bedtools intersect -u -a integration_site_annotation.txt -b gencode.v28.annotation_intron.bed >
SampleName_intron.bed
```

```
bedtools intersect -u -a integration_site_annotation.txt -b gencode.v28.annotation_intergenic.bed
> SampleName_intergenic.bed
```

iii. **Calculating genomic distance of vector integration site within 50,000bp upstream and downstream of TSS**

```
awk 'BEGIN{LIM=50000; FS=OFS="\t";}FNR==NR{tss[$1][$2]=1}FNR!=NR{for(i in tss[$1]){d=$2-i;
if(d>-LIM&&d<LIM) print $1,$2,i,d >> FILENAME".distance"}}'
<gencode.v28.annotation_gene_TSS.bed> < integration_site_annotation.txt >
```
